# Direct Sequencing of RNA with MinION Nanopore: Detecting Mutations based on Associations

**DOI:** 10.1101/575480

**Authors:** Noam Harel, Moran Meir, Uri Gophna, Adi Stern

**Affiliations:** School of Molecular Cell Biology and Biotechnology, Tel-Aviv University, Tel-Aviv, Israel

## Abstract

One of the key challenges in the field of genetics is the inference of haplotypes from next generation sequencing data. The MinION Oxford Nanopore sequencer allows sequencing long reads, with the potential of sequencing complete genes, and even complete genomes of viruses, in individual reads. However, MinION suffers from high error rates, rendering the detection of true variants difficult. Here we propose a new statistical approach named AssociVar, which differentiates between true mutations and sequencing errors from direct RNA/DNA sequencing using MinION. Our strategy relies on the assumption that sequencing errors will be dispersed randomly along sequencing reads, and hence will not be associated with each other, whereas real mutations will display a non-random pattern of association with other mutations. We demonstrate our approach using direct RNA sequencing data from evolved populations of the MS2 bacteriophage, whose small genome makes it ideal for MinION sequencing. AssociVar inferred several mutations in the phage genome, which were corroborated using parallel Illumina sequencing. This allowed us to reconstruct full genome viral haplotypes constituting different strains that were present in the sample. Our approach is applicable to long read sequencing data from any organism for accurate detection of *bona fide* mutations and inter-strain polymorphisms.

## Introduction

A major goal of genetics today is the characterization of genetic diversity in a population. In microbes, such diversity is generated in particular by high mutation rates, which may generate both nucleotide substitutions as well as point insertions or deletions (indels) (*1*). Longer indels may also occur (*2*), as may events of genetic recombination. The resulting genetic diversity has significant implications for epidemiology, molecular and serologic diagnosis, pathogenesis, and therapeutic management of disease (*3–9*). Thus, new molecular methods are being continuously developed to improve the characterization of high intra-host genetic diversity of microorganisms (*10–12*). The availability of second-generation DNA sequencing technologies (*13*), with the Illumina platform currently at the forefront, has made the sequencing of genomes conventional. In particular, this technology has dramatically furthered the study of viruses, whose relatively small genomes allow in depth characterization of a population of viruses (*14, 15*).

Illumina-based sequencing allows the detection of minor variants that the standard Sanger-based method often missed. However, Illumina short-read sequencing technologies all share one major limitation, the short length of each read, which typically ranges between 75 and 600 bp (for a paired end read). This means that a complete viral genome sequence cannot be obtained in a single read, impairing the ability to link distant mutations in an individual genome. Another Illumina limitation is that RNA cannot be sequenced directly. During library preparation, RNA is transcribed into cDNA and amplified by PCR. This creates multiple problems that have been extensively discussed but not resolved (*10*): first, reverse transcription and PCR may introduce errors during early stages of amplification that will be carried on to later stages (*16–18*). Second, some molecules may be preferentially amplified over others, a term known as PCR bias. Third, PCR and reverse transcription reactions can often result in chimeric DNA sequences that originate from different molecules (*19–21*). Together, these problems make the inference of haplotypes from PCR-based libraries that are sequenced with Illumina extremely limited.

Currently, single-molecule third-generation sequencing systems, such as Oxford Nanopore Technologies, provide a promising alternative for sequencing full-length single viral genomes (*22*). In fact, these technologies now allow directly sequencing either DNA or RNA. The long reads provided by these methods have the potential to allow for the inference of up to an entire genome of a typical RNA virus, whose genome is generally shorter than 10,000 bp (*23–27*). However, one of the major shortcomings of the third generation technologies are their relatively high error rates, with the proportion of errors on a read often exceeding 10% (*28, 29*). This high error rate makes the detection of true single-nucleotide variants very difficult (*7, 30, 31*).

Here, we devised a simple statistical procedure called AssociVar that allows weeding out real mutations from technical errors using *only* the MinION sequencing results. We focused our analysis on the MS2 bacteriophage, an extremely small (3,569 bases) and fast evolving +ssRNA virus (*32, 33*) that is highly amenable to direct sequencing with Oxford Nanopore MinION. We sequenced virus populations in parallel using both MinION and Illumina, allowing us to corroborate the inferences of AssociVar. This then allowed us to directly infer relationships between mutations and to deduce the entire genome sequences of viral strains in the population. We were also able to use AssociVar to analyze a yeast mRNA sample and data of mixed strains of Zika Virus. This illustrates the generality of our approach which can be applicable to other organisms as well.

## Materials & Methods

During the course of an evolve-and-resequence experiment, we performed serial passaging of the phage MS2 (Moran et al., in preparation). Briefly, clonal MS2 stock was propagated from a single plaque that was the precursor of all the evolutionary lines established in this work. We performed 15 serial passages at 37°C with two biological replicates (hereby denoted as A and B). The serial passages were performed as follows: 100 ml cultures of naive *E. coli* c-3000 were grown up to an optical density of OD_600_=0.4 (corresponding to a density of about 10^8^ cells/ml). Each passage was infected with 10ml of 10^9^ phages from the previous passage. The cultures were grown for 16 hours at each temperature with shaking, and the *E. coli* cells were then removed by centrifugation. The supernatant was subjected to filtration with 0.22 μm filter (Stericup^®^ Filter, EMD Millipore) to remove any remaining residues. Naive hosts were provided for each passage. The new phage stock was then stored at 4°C. Aliquots of these phage stocks were used for measuring the concentration of phages by plaque assay and infecting the next serial passage. We then determined the population frequency of each mutation at passage 1 and passage 15 through whole genome deep sequencing as described below, using Illumina and MinION.

### Illumina library preparation

Library preparation was performed according to our lab AccuNGS sequencing protocol (*34*) with some modifications: The reverse transcription reaction was performed using SuperScript^®^ III Reverse Transcriptase (Thermo Scientific), the reaction was performed on 500 ng of RNA phage with 1 μl of dNTP mix (10 mM), 1 μl of R4 primer (TGGGTGGTAACTAGCCAAGCAG) (10 μM) and 10 μl of sterile distilled water. The mixture was incubated for 10 minutes at 65°C followed by incubation on ice for 7 minutes. After brief centrifugation, 4 μl of 5X First-Strand Buffer were added along with 1 μl DTT (0.1 M), 1 μl of RNase OUT (Thermo Scientific) and 1 μl of SuperScript™ III RT (200 units/μl). The mixture was incubated for 60 minutes at 55°C followed by inactivation for 15 minutes at 70°C. The last step was adding 1 μl of RNase H to the reaction according to manufacturer instructions. cDNA from the reverse transcription reaction was directly used as a template for the PCR amplification of the full MS2 genome in three overlapping fragments. PCR reactions were performed using “Phusion High-Fidelity DNA-polymerase” (Thermo Scientific) according to manufacturer instructions with the primers: 1F (GGGTGGGACCCCTTTCGG), 1R (TTTTTCTAGAGAGCCGTTGCCT), 2F (GGCCCAAATCTCAGCCATGC), 2R (CGTGTCTGATCCACGGC), 3F (GGCACAAGTTGCAGGATGCA), 3R (TGGGTGGTAACTAGCCAAGCAG). After PCR, the three amplicons were purified with a PCR clean up kit (Promega). Purified amplicons were quantified with Qubit assays (Q33216, Life Technologies), diluted and pooled in equimolar concentrations. The Illumina Nextera XT library preparation protocol and kit were used to produce DNA libraries, according to manufacturer instructions with some modifications. Briefly, the tagmentation reaction was performed with 0.8 ng/μl of DNA, 10 μl TD Buffer, 5 μl ATM enzyme and up to 5 μl DDW. The mixture was incubated for 5 minutes at 55°C and then directly used as a template for 50 μl PCR reaction using “Phusion High-Fidelity DNA-polymerase” (Thermo Scientific) according to manufacturer instructions with the index primers from the Nextera XT kit. After PCR, a double sized selection was performed using Ampure beads to remove short and long library fragments, since 350bp fragments were required. We collected the supernatant and read 2 μl from each sample in the TapeStation (Agilent high sensitives D1000) to verify fragment size. The yeast enolase sample was prepared as described for the phage RNA. RT and PCR amplification were performed using primers: E1 (ATGGCTGTCTCTAAAGTTTACGCTA) and E2 (TTACAACTTGTCACCGTGGTGG). Libraries were sequenced on an Illumina Miseq using the 2X250 MiSeq reagent kit (Illumina, MS-102- 2003) for paired-end reads.

### Illumina MiSeq Read mapping

Bioinformatics processing of the data was performed using the AccuNGS pipeline (*34*) with the default parameters (minimal %ID=85, e-value threshold= 1E-07 and q score cutoff of 30). Briefly, this pipeline is based on (a) mapping the reads to the reference genome using BLAST, (b) searching for variants that appear on both overlapping reads, (c) calling variants with a given Q-score threshold and inferring their frequency. All libraries attained a mean coverage of ~10,000 reads/base. The reference genome was determined by comparing the consensus of passage 1 to GenBank ID V00642.1 (differences noted in Table S1). When examining the results of the control sequence (a plasmid bearing the MS2 genome), we noted a high error rate at several positions that resided near the primer sites, and accordingly 30 positions from each end of the genome were excluded from downstream analysis.

### Long-read Oxford Nanopore MinION sequencing

The Oxford Nanopore MinION was used to sequence the MS2 RNA directly. We sequenced the three samples (p15A, p15B and p1A) in three separate runs. We prepared direct RNA libraries according to manufacturer library prep instructions with some modifications. We altered the supplied reverse transcriptase adapter (RTA) (*22*), which has a T10 overhang, to specifically target the MS2 genome with 23 nucleotides complementary to the MS2 conserved 3’ end. The ligation reaction was performed with 500 ng RNA in 9 μl, 100 nM costume adapter in 1 μl, 10 μl of NEBNext Quick Ligation Buffer and 1.5 μl of T4 DNA Ligase enzyme (NEB). The mixture was incubated for 10 minutes at 65°C followed by incubation on ice for 2 minutes and then directly used as a template for the next step of the library prep which is cDNA synthesis according to manufacturer instructions. The cDNA synthesis step was performed in order to maintain the RNA fragments integrity during MinION library prep and sequencing. The library was cleaned up each time using 2 μl of AMPure XP DNA beads per 1 μl of sample and we added 3 μl of RNase OUT (Thermo Scientific) to protect the RNA. The RNA was directly sequenced on the MinION nanopore sequencing device using a FLO-MIN107 flow cell equipped with the R9.5 chemistry. The MinKNOW control software version 1.14.1 was used and was allowed to proceed for 48 hours. The basecalling was performed locally by the Miniknow software as well, and the data was written out in the FASTQ format. Reads were filtered with the MiniKNOW default cutoff of a minimum average qscore of 7. The AccuNGS pipeline (*34*) described above was next applied to the data in order to determine variant frequencies. BLAST parameters were modified to minimal % identity=60, e-value threshold= 1E-07 and no q-score cutoff, to allow the highly variable minion reads to map. As with the Illumina results, 30 positions from each end of the genome were excluded from downstream analysis. This also solves the problem of very low coverage areas in the MinION sequencing.

### MinION Variant Distributions and Error Rate Estimation

Variant frequency distributions for MinION and MiSeq were calculated by using the AccuNGS pipeline results. For substitutions and deletions, the distribution is straightforward and accounts for the results for all bases. For insertions, we focused on point insertions, defined as the first insertion after any position, in order to create the frequency distributions. Positions close to the ends of the genome that show a high error rate in MiSeq due to primer proximity were removed from this analysis. To calculate the error rate, we sequenced a control sample from the beginning of the passaging experiment (p1A), and the mean, 95^th^ percentile and 99^th^ percentile errors were calculated for every error type.

### AssociVar: Association Tests to Identify Real Mutations

AssociVar searches for strong associations between variants as an indication that these represent *bona fide* mutations. The method is based on five stages:

a. <u>Detecting non-random associations</u>. For each pair of positions, a read is classified into four categories based on whether the read bears the WT nucleotide (i.e., identical to the refence genome) or non-WT nucleotide (i.e., different from the refence genome) at each of the two given positions. We use this to create a 2×2 contingency matrix of observed frequencies, which is then used as the input for a chi-square test and a resulting chi-square statistic. We focus only contiguous reads that spanned the entire genome. Notably at this stage we focus only on WT versus non-WT assignments (rather than the exact identity of the non-WT allele) for computational tractability. This is relaxed later on.
b. <u>Removing proximal positions</u>. Since we observed that positions that are highly proximal (< 15 bp apart) often tended to be highly associated, and we suspected this is an artifact of the sequencing machine, chi square results for all such proximal positions were removed from the analysis.
c. <u>Normalization of the chi-square statistic</u>. To make the different positions comparable, we normalized the chi square statistics per position by calculating a modified z-test score *z* for the chi-square statistic *x* of each pair of positions (*p*_1_, *p*_2_). This was done by dividing the difference between *x* and the median by the median absolute deviation (*MAD*) and multiplying the result by 0.6745, as follows: *z* = 0.6745 ∗ (*x* − *median*)/*MAD*. Thus, for each position *p*_1_ we calculated its median and its median absolute deviation across the chi-square statistics for position *p*_1_ and any other position *q*. This essentially served to test whether some positions display strong outliers in their chi-square statistics (*35*), as can be seen visually for *bona fide* mutations displayed in Figures 5 and 6.
d. <u>Local maximum analysis</u>. Because positions proximal to each other tend to present spurious high associations, due to transitivity we expect a position *next* to a real mutation to also be highly associated with other positions with real mutations. In other words, if positions *p* and *q* are highly associated because they are real mutations, and positions *q* and *q* + 1 are highly associated because they are proximal, we will see a high association between positions *p* and *q* + 1 as well. However, we expect the association between the real mutations to be the highest, i.e., to be a local maximum in the surroundings of a given pair. A normalized chi score’s surroundings is defined as the four neighboring normalized chi scores when the data is regarded as a two-dimensional matrix. For example, the normalized chi score for (*p*, *q*) is required to be higher than the normalized chi scores for (*p*, *q* − 1), (*p, q* + 1), (*p* − 1, *q*) and (*p* + 1, *q*).
e. <u>Use of a control sequence</u>. In order to create a cutoff for the normalized chi-square statistic, we used the values obtained for a control sequence (Fig. S3). We know that our control sample was not completely homogenous, since it contained two mutations at a frequency > 10%. Nevertheless, it served as a valid control when setting a confidence rate of 99.9%, i.e., calculating the normalized chi score that allows 0.1% of the positions in our control sample to be identified as significant (allowing for 3 “false positive” positions in our case).

After stages (a) through (e), AssociVar infers *n* positions with real mutations in the population. The last stage is to identify the identity of the mutations (A, C, G, T, −), in a similar way to that described in (a). Insertions are ignored here. Every position has four possible alternative variants (the three nucleotides that differ from the reference, or a deletion), and we test these variations against each other using chi-square tests, leading to 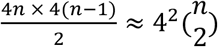 different tests, where *n* is the number of positions previously identified as having real mutations. Again, we created a 2×2 contingency matrix of observed frequencies, which is then used as the input for the chi-square test. For every position, we choose the variant with the highest average chi-square statistic for all the tests for pairs containing that variant.

### Haplotype/Strain Identification Analysis

We begin by focusing only on the variants inferred as bona fide mutations in the last stage. This means that in principle there are 2^*n*^ possible haplotypes bearing these mutations. We filter out reads with variants that do not match our inference (for example, if one of the inferred mutants is A535G, we filter out reads with the nucleotides C, T or a deletion at position 535). We then use our inferred 95 percentile error thresholds (Table 1) to deduce which combinations of mutations are likely to be true and which may have been created by the technical error rate.

**Table 1.** Estimated error frequencies for MinION sequencing. Shown are mean frequency values, 95^th^ and 99^th^ percentile values for each type of error, based on the control sample (Methods).

|  | Mean Error Frequency | 95<sup>th</sup> percentile Error Frequency | 99<sup>th</sup> percentile Error Frequency |
| --- | --- | --- | --- |
| Substitutions | 0.07 | 0.21 | 0.33 |
| Deletions | 0.07 | 0.24 | 0.43 |
| Insertions | 0.01 | 0.04 | 0.08 |

We use an iterative approach to classify which of the 2^*n*^ haplotypes is reliable. First, for every single nucleotide variant, we group together all of the haplotypes that include this base variant (a haplotype can appear in more than one group). Second, in each group, we compute the relative frequency of each haplotype as its proportion of all haplotypes in the group. We iterate through the haplotypes from highest frequency to lowest, classifying each haplotype as reliable or not. The first haplotype is automatically classified as reliable. For every haplotype, we compare its relative frequency with the probability that it is created by technical errors from the closest haplotype classified as reliable, called its parent haplotype, using the inferred error threshold. For example, if a haplotype has an additional deletion and substitution when compared to its parent haplotype, we require that its relative frequency be higher than the product of 0.214×0.237=0.051 to be classified as reliable (using the 95^th^ percentile error frequencies in Table 1). We iterate through the haplotypes until classifying all the haplotypes in each group. If reads for a WT haplotype exist, the WT is also treated as a base for a group – which in this case will include all of the observed haplotypes. Haplotypes may appear in more than one group, if a haplotype appears as reliable in at least one group it will be classified as reliable overall. See supplementary text for a visual summary of the algorithm we employ.

The code we provide produces a file with all the variant combinations observed, the calculations described here and whether a haplotype is reliable or not. It also produces a file with the haplotypes classified as reliable, with their proportion in the population recalculated appropriately.

### Sequencing Data and Code Availability

The sequencing data created and used in this study is available in the Sequencing Read Archive (SRA, https://www.ncbi.nlm.nih.gov/sra), under BioProject PRJNA547685 (https://dataview.ncbi.nlm.nih.gov/object/PRJNA547685?reviewer=vef29oif04j9mpj8229qglj1bf)

The accompanying code can be found at https://github.com/SternLabTAU/AssociVar.

## Results

We set out to sequence two evolved populations of the MS2 coliphage. Both populations were derived by fifteen serial passages performed at 37°C (denoted as A and B) as part of an evolve-and-resequence experiment (Methods). We first performed deep sequencing of both populations at passage 15 with the Illumina MiSeq platform (*34*). This revealed several segregating mutations (Fig. 1), some of which shared similar frequencies. However, due to the short-read nature of the sequencing it was impossible to infer whether these mutations co-occurred on the same genome.

**Figure 1.**
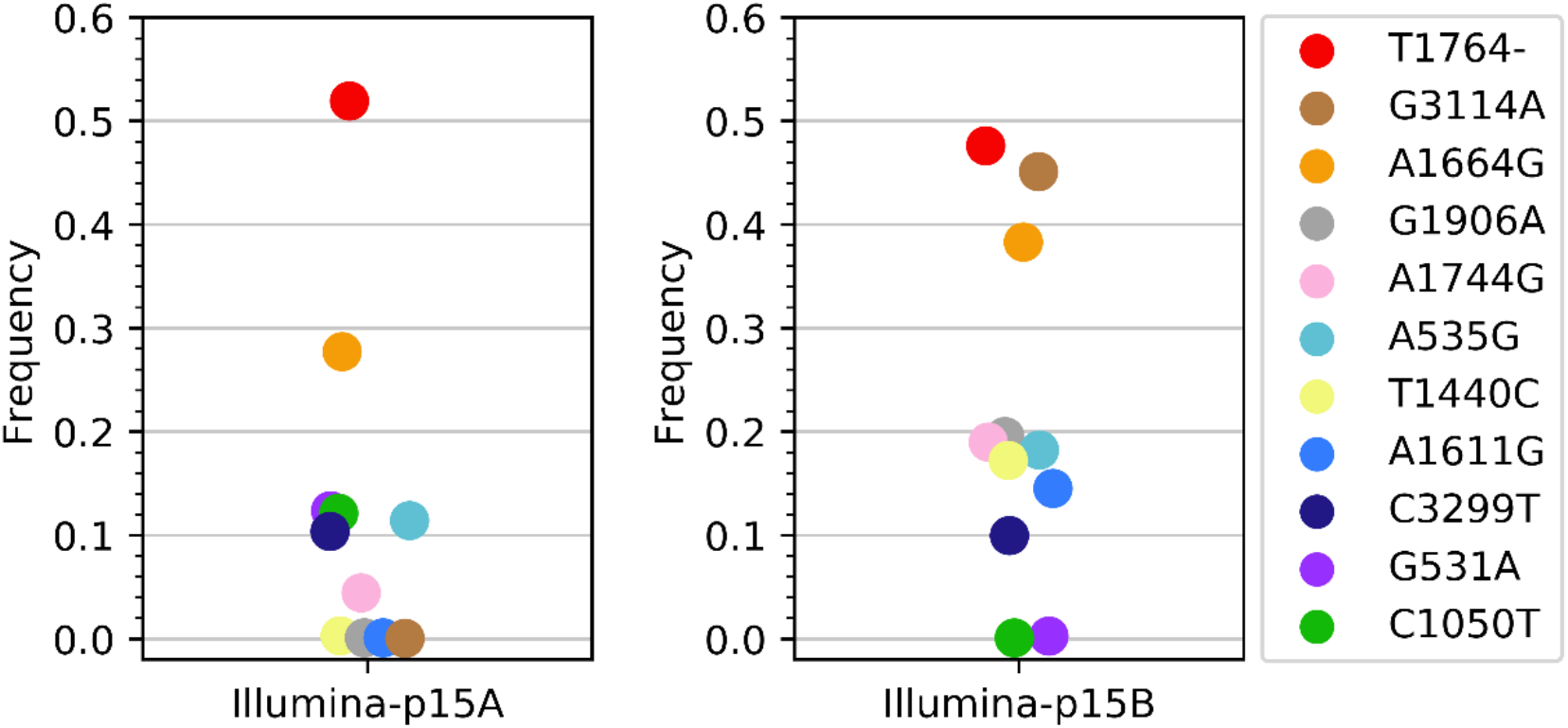
Mutation frequencies in two MS2 phage populations evolved over 15 passages, based on Illumina sequencing. Shown are the frequencies of mutations that were detected at a frequency above 10% in passage 15 in either of the two replicate populations. Mutations are spread horizontally for visual clarity only.

We next sequenced the same two populations of RNA viruses from passage 15 using Oxford Nanopore’s MinION. Importantly, we employed direct RNA sequencing, without using reverse transcription or PCR amplification, and without any shearing of the genomes. The only requirement for library preparation was the ligation of an adaptor to the 3’ of the RNA genome, allowing the 3’ to enter the sequencing pore. Each replica was sequenced independently, denoted as p15A and p15B. We also used MinION to sequence a sample from line A passage 1. As this was a mostly unevolved and homogenous population, we used this as a control sample.

### Read Lengths and Alignment

A total of 417,000, 105,000 and 400,000 reads were produced for the MinION-p15A, MinION-p15B and the control runs, respectively. In order to map the reads to the MS2 reference genome we ran our computational pipeline (Methods) (*34*), which infers the proportion of each point mutation (A, C, G, T, or “−“) at each position in the genome. Over 97% of the reads were mapped to the reference, yet often sequencing terminated before it reached the 5’ end in both the evolved and control populations (Fig. S1). Nevertheless, approximately 15% of the reads (between ~15,000 and ~60,000) covered the entire MS2 genome.

### Distribution of observed variants and MinION error rates

We next focused on the frequency of an observed variant, defined here as any base called differently from what is present in the reference sequence, at any position. We expect such variants to be the sum of two independent processes: real biological variations derived from evolutionary processes in the phage populations, and technical errors introduced by the sequencing process. Comparing between the variant distributions of Illumina and MiniON, it was evident that MiniON suffers from a very high technical error rate (Fig. 2). Notably, in the control population the number of variants exceeding a frequency of 1% in the Illumina sequencing was 12, whereas with MinION we observed 8917 variants in 3326 positions exceeding 1%. This allowed us to infer that the vast majority of MinION variant frequencies are technical errors, and further allowed us to roughly estimate the various types of MinION error rates for our experiment (Table 1). Notably, we observed that the point deletion and point insertion rates together exceeded the substitution rate, reinforcing previous observations (*36, 37*).

**Figure 2.**
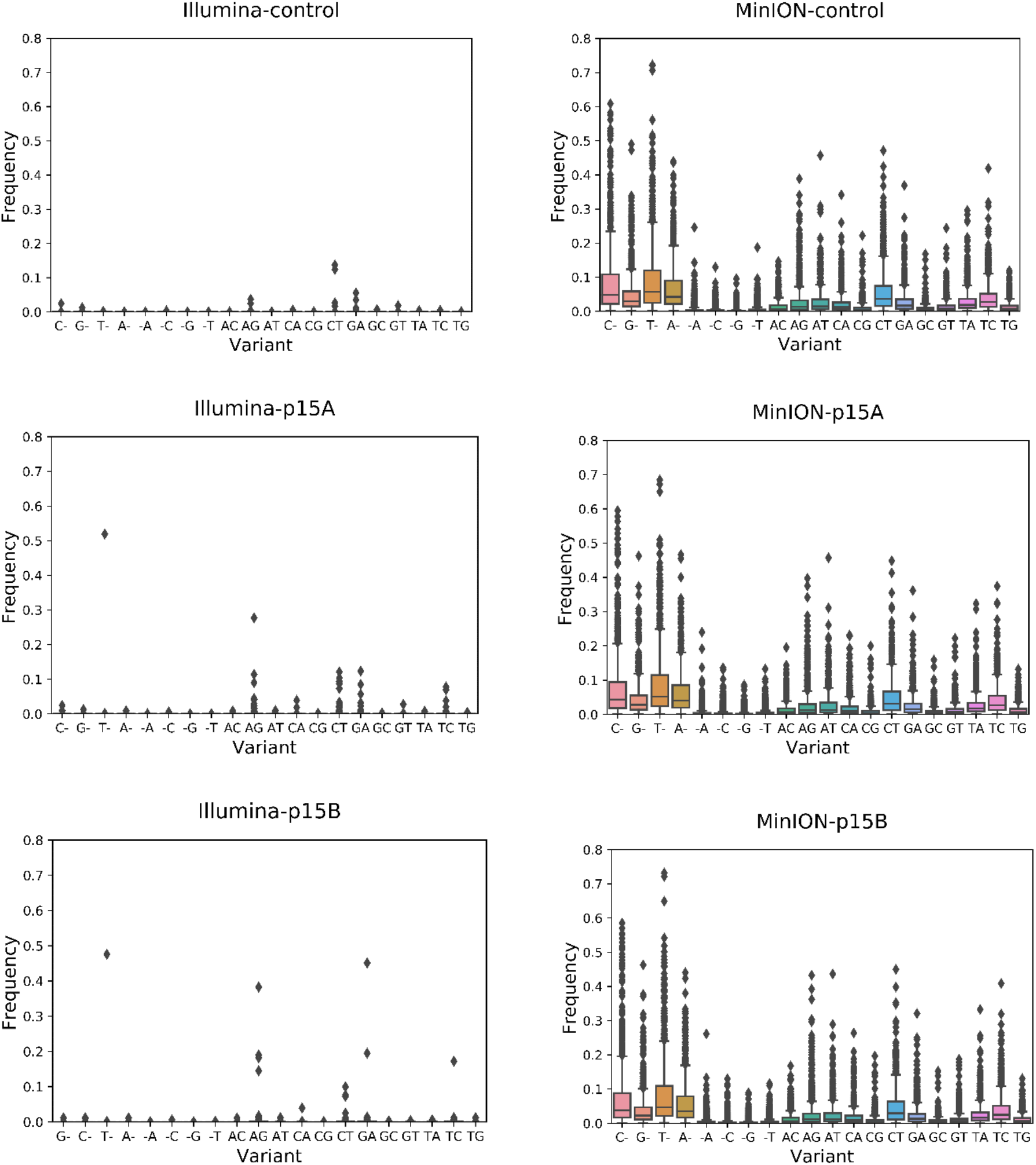
Variant distributions from the populations sequenced using Illumina (left) and MinION (right) for samples control, p15A and p15B. Variants are broken down by the type of mutation they cause, substitutions or point deletions. “X-“indicates a single base deletion, “-X” indicates a single base insertion and “XY” indicates X replaced by Y (where X and Y are any of the four bases).

We attempted to use the inferred MinION error rates as thresholds that can distinguish between real mutations from errors, by setting the 95^th^ percentile obtained for the control sample as an error threshold for each type of error (Table 1). This naïve approach that is often used, quickly turned out to be invalid, as corroborated by our parallel Illumina results. For example, we knew from the Illumina results that only 6 mutations in line A and 8 mutations in line B exceeded a frequency of 10% at passage 15 (Fig. 1). However, the MinION results showed 1168 mutations in 949 positions and 1081 mutations in 879 positions exceeding 10% in both replicas, respectively.

### Associations between variants in MinION

We sought a strategy to weed out the technical errors from the real mutations in the MinION results independently of the Illumina results. We calculated the conditional probabilities of observing one variant given another variant observed on the same read. When observing the pattern of conditional probabilities (Fig. 3), we noted two distinctly different patterns. Some variants co-occurred more or less randomly with all other variants, manifested as more or less the same probability of observing one variant given any other variant (similar colors across a given column in Fig. 3A). On the other hand, some variants displayed a non-random pattern, where the probability of observing variants together depended very much on which pair of variants was examined (different colors across any given column in Fig. 3B).

**Figure 3.**
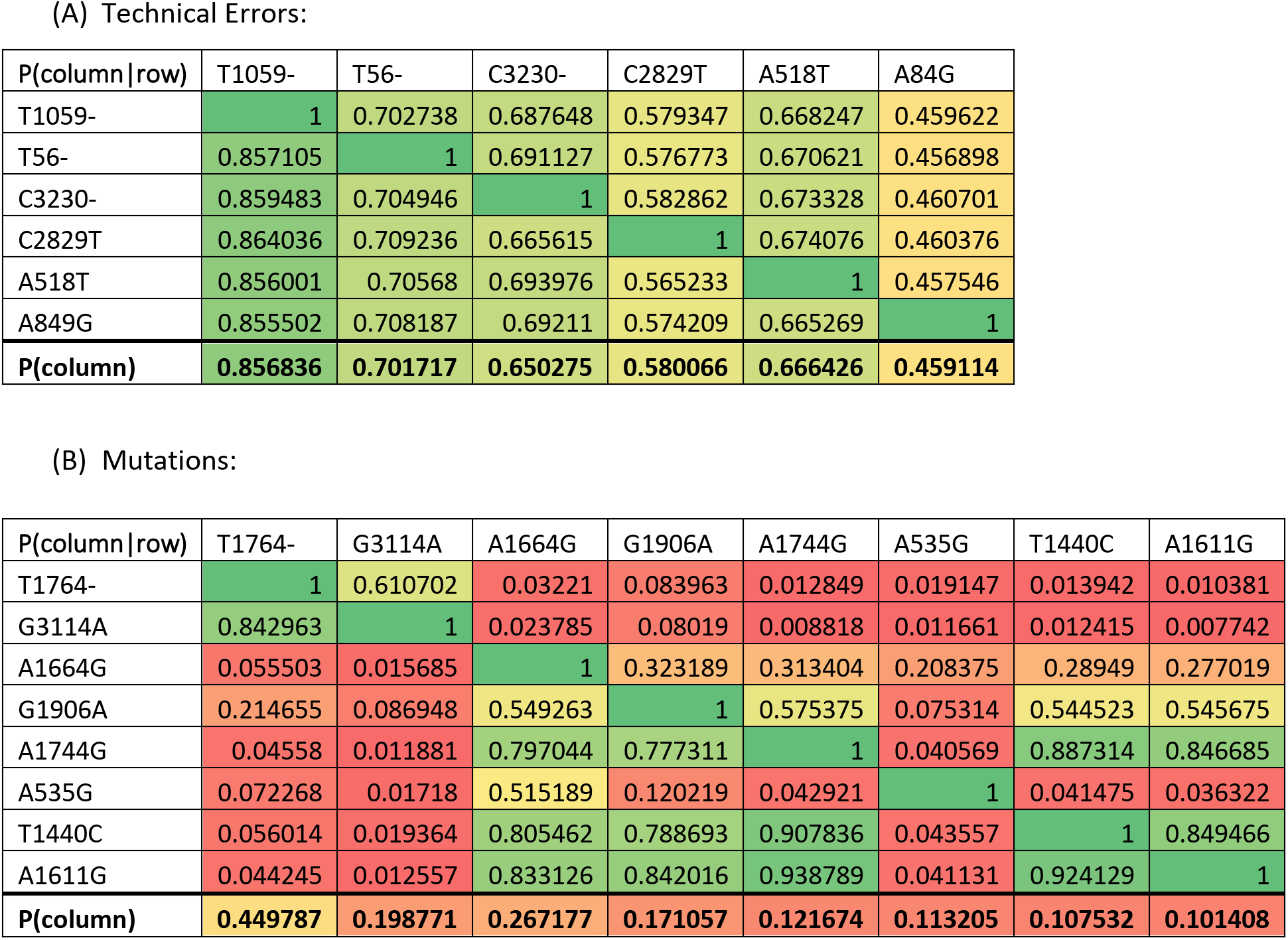
Conditional probabilities P(column|row) of observing one variant (column) in the presence of a second variant (row) on the same read. Bottom row shows P(column), probability of variant in column. Cells are color-coded using a gradient to highlight similar values across cells. (A) The highest frequency-presenting variants in MinION for line B (top three substitutions and top three deletions), all of which were *not* detected using Illumina and thus classified as technical errors. Values across a given column are more or less identical (with the exception of seeing a variant with itself), showing that P(X|Y) = P(X). (B) The mutations identified by Illumina in line B. Values across a given column are sometimes dramatically different. For example, looking at the A1744G column, A1744G tends to be highly associated with T1440C and A1611G, somewhat associated with A1664G and G1906A, and not at all associated with A535G, T1764- and G3114A. This pattern supports the fact that these are all *bona fide* mutations rather than technical errors.

Importantly, the variants that displayed a non-random pattern were variants that we knew were true mutations based on the Illumina data. This led us to realize that random technical errors are expected to display a different pattern than real biological mutations: we expect technical errors to be associated randomly with *any* other technical error, whereas a pair or more of real biological mutations are expected to be non-randomly associated with each other. This is a reflection of evolutionary processes operating on genomes. While mutations created from replicative polymerases will be mostly randomly distributed along the genome, selection and genetic drift will lead to the fact that specific combinations of mutations reach higher frequency. Thus, true mutations that are prevalent in a population will tend to be either present with some other mutations on the same genome/read, or *not* present with some other mutation on the same genome. Both these properties (tendency to be present or not present with other mutations) reflect non-random association between mutations.

One of the most commonly used methods to test for associations between two properties is the chi-square test: here we use this test to see whether the observed joint variant counts deviate from what is expected when variants are counted independently. To this end, each variant was classified as either wild-type (WT) or non-WT, based on whether it was identical or not to the reference genome. Notably, this led to 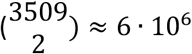 different tests. Because the observation count highly affects the chi-square result and the MinION sequencing coverage increases greatly along the genome, we only used reads spanning the entire genome of the virus for this analysis (Methods).

We began by inspecting all associations between all pairs of positions. This allowed us to make a few general observations. First, we observed that proximal pairs of positions (residing up to 15 bases apart from each other) tended to be highly associated. We postulate that this is a reflection of MinION errors, and also the high deletion rate, which could cause slight misalignment of reads covering positions proximal to the deletions. Second, we observed a very similar pattern of associations among the three samples. This suggests that MinION sequencing has a tendency towards specific a pattern of errors for a given genome that is sequenced (Fig. S2).

We devised a method called AssociVar that detects the real variants in the data, based on the following properties: (a) the method searches for the strongest non-random associations, (b) the method takes into account that pairs of proximal positions (i.e., up to 15 bases apart) that have high associations between them are likely false positives induced by the MinION machine itself, (c) in order to make the different positions comparable, the method normalizes the distribution of chi square scores per position, essentially searching for outliers from all the associations of a given position, (d) the method also takes into account that because proximal positions are highly associated, a position next to a real mutation may be associated with other real mutations due to transitivity. However, we expect the two real mutations to display the highest association, i.e., we require an association to be a local maximum in a given window. Finally, (e) the method uses a control sample to set a cutoff for the highest associations (Methods). AssociVar hence calculates a normalized chi-square statistic and infers the positions where “true” variation occurs, based on the above properties (Fig. 4).

**Figure 4.**
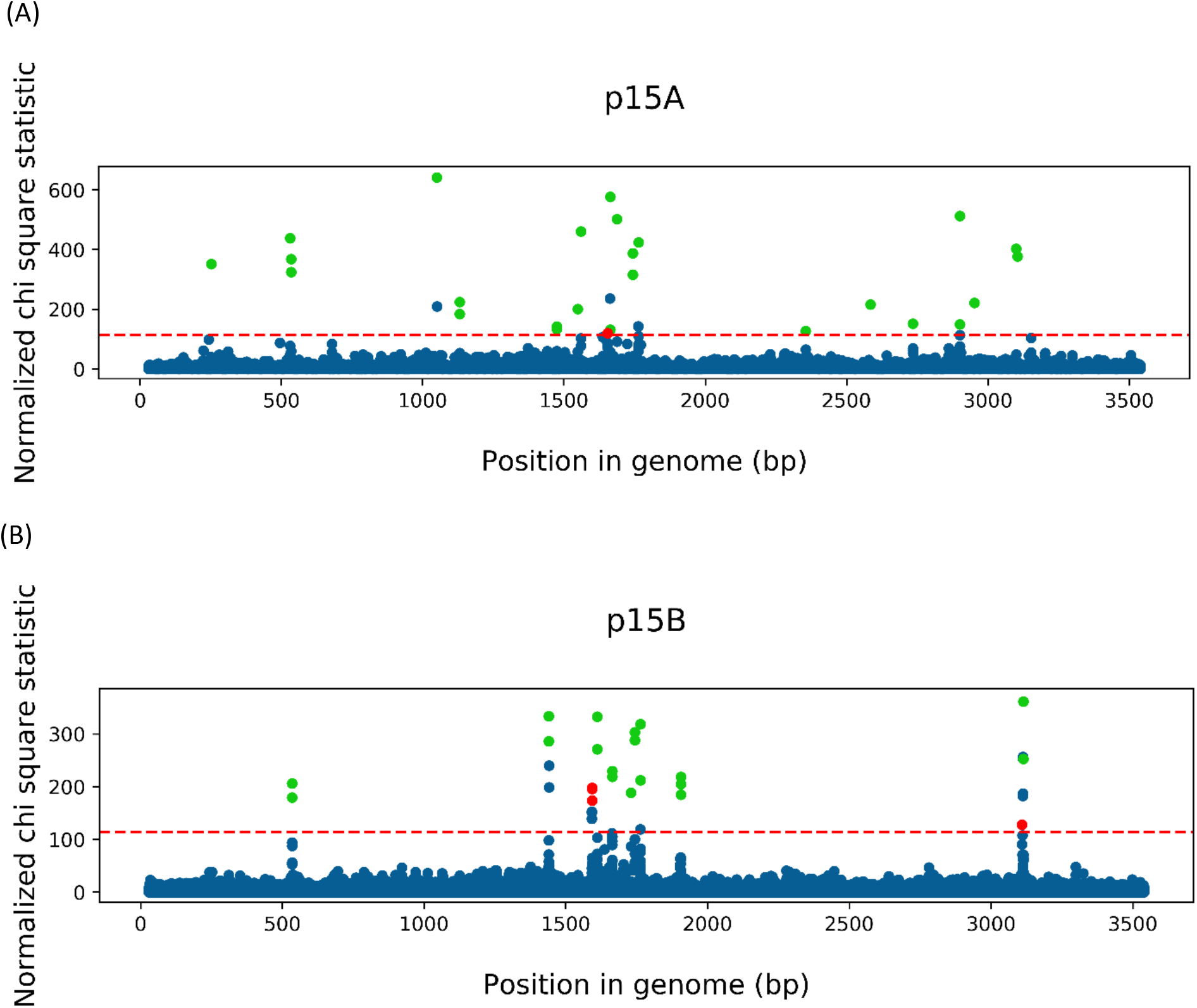
Chi-square statistics plotted along the genome for the (A) p15A and (B) p15B MinION sequencing. Each dot represents the normalized chi-square statistic between a given variant and another variant at a position more than 15 bp apart. Associations marked in blue are classified as non-significant. Associations marked as green are true positives, where the position in the genome they identify is a real mutation as verified by Illumina sequencing. Associations marked as red identify positions that are false positives and do not have real mutations according to the Illumina results (one position for p15A and two positions for p15B). The red line illustrates the cutoff defined by the control sequence. Notably, the blue dots above the line are positions *near* real mutations, removed by AssociVar via the requirement for a local maximum (Methods).

After applying AssociVar to the data, we were able to identify five out of the six mutations appearing at a frequency above 10% in the Illumina results in p15A, and all eight positions within the p15B sample (Fig. 4, Table S2). Notably, AssociVar also often correctly identified mutations segregating at lower frequencies (1-10%) according to Illumina. When focusing on the false positive rate of the method, AssociVar reported one out of 3,467 positions as false positives for p15A and two out of 3,475 for p15B, where false positives are defined as positions identified in the association analysis but segregating at a frequency lower than 1% according to Illumina. All in all, the results indicate that our association approach has the power to resolve real variants from technical errors based on the MinION data alone.

We began the analysis by classifying mutations into WT and non-WT, for computational tractability. Next, identifying the specific *nucleotide* variant in our samples after having identified the positions with real mutations is easy enough using a similar approach. Every position has four possible variants (the three nucleotides different that the reference and a deletion for that base), and we test these variations against each other, again – under the assumption that the most highly associated variant for each position is the real one (Methods). For the positions identified by our association analysis and verified as correct by the Illumina sequencing, all but one of the positions were matched with their correct variant with this method (position 3114 in p15B was identified as a deletion instead of nucleotide A).

### Identifying mutations in the absence of a control sample

The MinION RNA sequencing kit comes with a control sequence of the enolase II yeast gene. We ran our association analysis on the enolase results, originally to verify that we pick up no variation in this gene. However, we were surprised to see two positions with a very high and outstanding association (Fig. 5).

**Figure 5.**
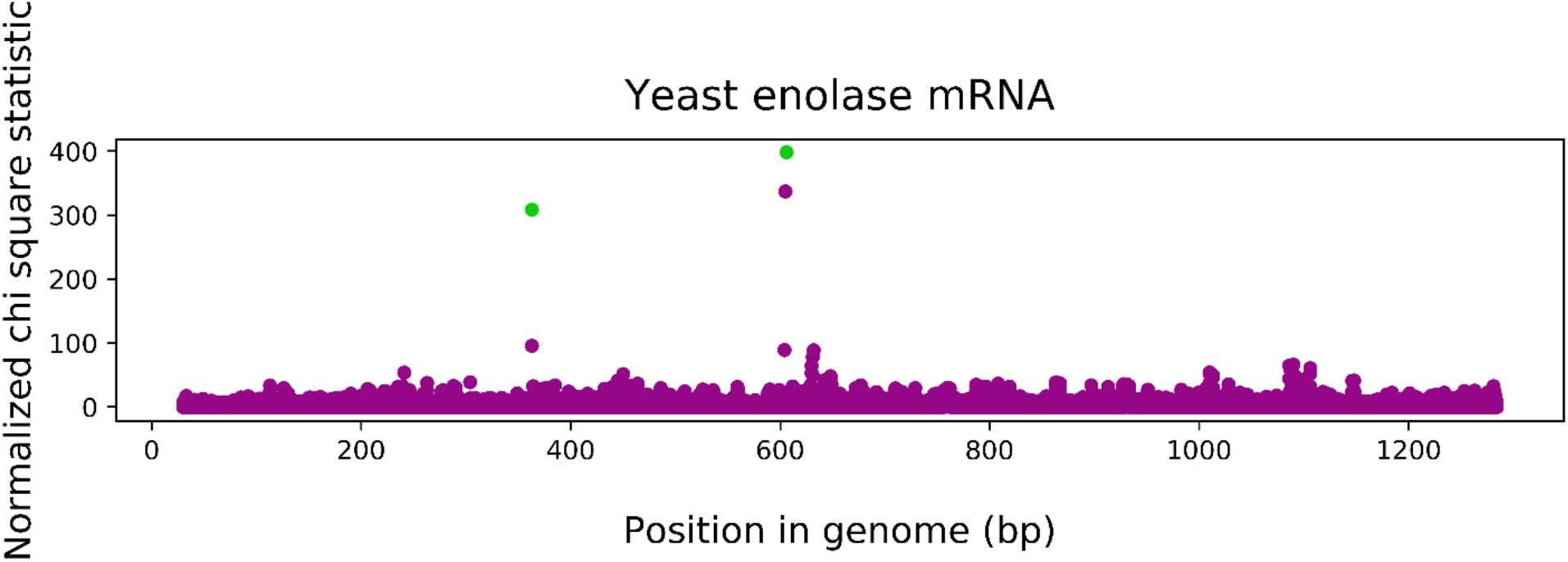
Chi-square statistics plotted along the gene for the enolase yeast gene. Two positions stand out as having a high association with each other, 363 and 606, evident by the two high peaks in the figure (green). Notably, other high associations were ruled out as induced by proximity with the local maximum analysis (Methods). Since we do not have a control sample for this gene, we cannot use it to infer a cutoff. However, the association between these two positions is so prominent when compared to the rest of the data that we were able to conclude they are between positions with “true” variation (as later verified by Illumina).

We thus sequenced the same sample with the Illumina MiSeq platform. Reassuringly, the results verified the findings of AssociVar and showed that these two variants do appear in the sample and are the only two variants that appear there at a frequency higher than 10%. This suggests that our method can be used (a) as a general approach and not only for virus populations, and (b) in the absence of a control.

We next tested our method on a sample of Zika virus genomes (*30*). In this study, two different strains that differed at several positions had been artificially mixed and sequenced using MinION. We ran AssociVar on one of the sequenced amplicons, in the absence of a control sequence. At least five of the six true mutations in this amplicon stood out as having highly prominent associations (Fig S4).

Finally, we tested how our method fares in the absence of a control for our MS2 data, and compared the true positive rate versus false positive rate as a function of (a) thresholds set for the normalized chi-square statistic, and (b) a frequency threshold from the Illumina results that determine when we define a mutation as true or false. We further compared this to the use of a “naïve” approach where we use a varying frequency threshold for the MinION results (as described above). Our results show that AssociVar inference is consistenly much more accurate that the naïve approach, and moreover, can be used even to detect mutations at a frequency of 1%, at the risk of some false positives (Fig. 6).

**Figure 6.**
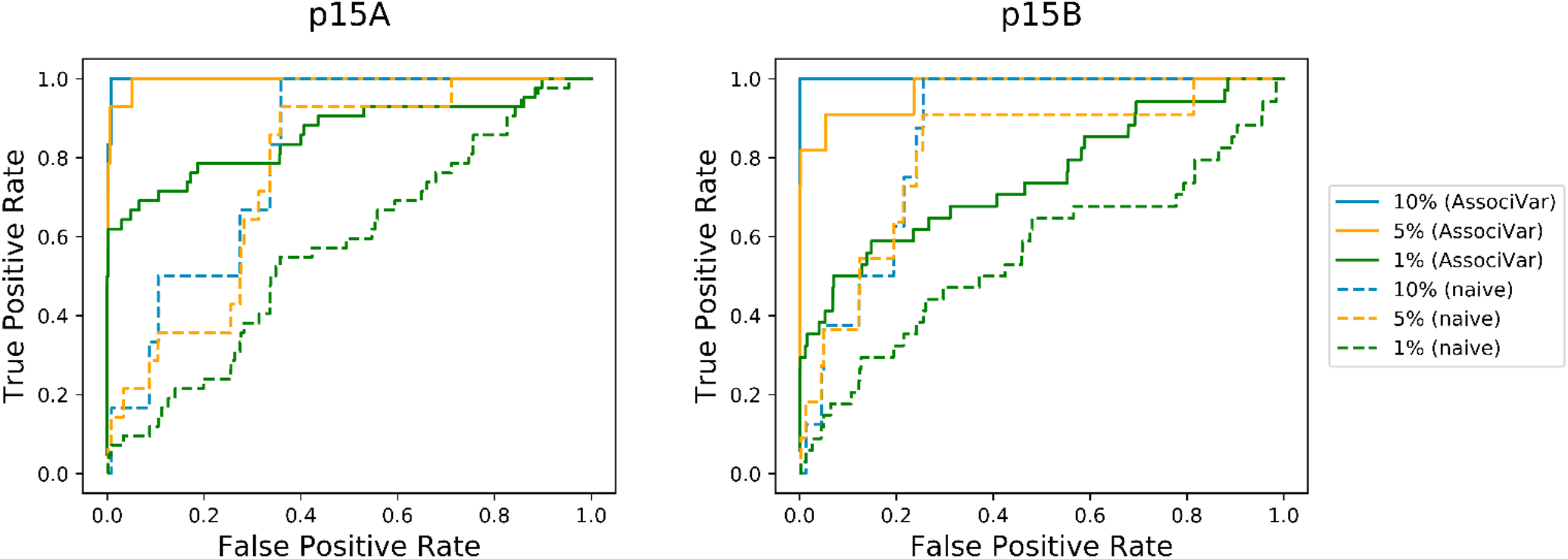
Receiver operating characteristic (ROC) of AssociVar versus a naïve method. Performance of prediction of mutations is assessed using ROC curves, where each curve is plotted as a function of the normalized chi-square statistic threshold for AssociVar (solid lines), or frequency threshold for the naïve method (dashed lines). The Illumina results are used as the gold standard test to define a mutation as true or false, and the three different colors represent different thresholds for this definition. For example, for the blue line labeled as 10%, only mutations at a frequency higher than 10% according to Illumina are defined as true.

### Haplotype/strain identification

One of the main goals of MinION sequencing, in particular in the context of RNA virus evolutionary experiments, is the detection of haplotypes and identification of distinct strains in the population. We thus set out to use the approach we devised to infer the composition of strains in our MS2 samples. Notably, this is challenging on two fronts: first, our association approach can tell us which mutations in the MinION data are real, and which pairs are associated, but it does not tell us what their frequency is (or rather, we do not trust the observed frequencies given the very high error rate). Second, we are interested in inferring haplotypes, i.e., which mutations reside together regularly on the same genomes and which do not. Once again, the high error rate makes this extremely tricky since we observe reads bearing almost all possible pairs of mutations. In fact, most reads bore so many variants, that around 15-20% of the bases called per read were different than the reference (Fig. S5). In this case, we used a two-pronged approach: we first focused only on variants inferred as true mutations using our method, AssociVar, as described above. Second, we used the inferred error threshold to infer the probability of two or more variants residing erroneously on the same genome, utilizing an iterative approach in which we compare a given haplotype to haplotypes already classified as reliable (Methods). In our case, because the MS2 populations bore many mutations at low frequencies, we also limited the analysis to variants that appeared at a frequency of at least 5% in the MinION sequencing results.

The summary of inferred strains is shown in Table 2 and provides a few interesting insights into our populations. First, twelve and sixteen different strains were identified in each of the A and B populations, respectively. Second, A1664G and T1764-, both of which rose to high frequencies in both replicates, were found to be mutually exclusive in both replicates. On the other hand, G3114- (which we know from the Illumina data to actually be G3114A) was found to be tightly linked to T1764-, in line with the very similar frequencies of these mutations in p15B. All of these results were unobtainable with the Illumina results alone and highlight the added value of using MinION for inferring viral genotypes.

**Table 2.** Strains in the p15A and p15B populations inferred by AssociVar and their associated read counts and frequency in the populations.

| Strain | Read count | Frequency |
| --- | --- | --- |
| **p15A** |  |  |
| T1764- | 3301 | 0.261 |
| A1664G | 2353 | 0.186 |
| T1764-, A2356G | 956 | 0.076 |
| C1724T, T1764- | 878 | 0.07 |
| C1131T | 784 | 0.062 |
| A535G, A1664G | 702 | 0.056 |
| T1475C, T1764-, T2953C | 589 | 0.047 |
| G531A, T1764-, C3100T | 578 | 0.046 |
| C1050T, T1764-, G2901A | 524 | 0.041 |
| A535G | 519 | 0.041 |
| T2953C | 431 | 0.034 |
| T1764-, G2901A | 387 | 0.031 |
| C1050T | 184 | 0.015 |
| T1475C | 168 | 0.013 |
| C3100T | 147 | 0.012 |
| G531A | 127 | 0.01 |
| **p15B** |  |  |
| T1764-, G3114- | 1203 | 0.307 |
| A1664G | 1006 | 0.257 |
| A535G, A1664G | 447 | 0.114 |
| A535G | 382 | 0.097 |
| T1440C, A1611G, A1664G, A1744G, G1906A | 309 | 0.079 |
| T1764-, A3109-, G3114- | 199 | 0.051 |
| T1440C, T1593C, A1611G, A1664G, A1744G, G1906A | 193 | 0.049 |
| G1906A | 106 | 0.027 |
| T1593C | 43 | 0.011 |
| A3109- | 21 | 0.005 |
| T1440C, A1744G | 9 | 0.002 |
| T1593C, A1611G, A1664G, A1744G, A3109-, G3114- | 1 | 0 |

## Discussion

We have developed here a simple and intuitive approach, AssociVar, to (a) detect *bona fide* mutations from MinION population sequencing, and (b) infer the set of haplotypes (strains) present in a population. Our approach is based on the notion that sequencing errors will be randomly dispersed along the reads, whereas real mutations tend to associate with specific genetic backgrounds. In the case where technical error rates are high (such as occurs with MinION), this allows one to focus on the real genetic diversity that is hidden in the vast array of technical errors generated by this method. Notably, our approach is general enough so that it can be used for any type of long read sequencing.

We applied AssociVar to sequencing data from an evolved population of phages where Illumina sequencing was available, allowing us to corroborate whether mutations we found based on analysis of the MinION data alone were indeed real. Strikingly, all but one of the high frequency mutations observed in the p15A and p15B data (>10%) were picked up using AssociVar, despite the fact that the 99^th^ percentile for technical errors was as high as 43% (Table 1). In fact, despite the very high deletion rate, AssociVar accurately identified the one real deletion mutation present in our populations, suggesting a very high sensitivity of the method. Our approach also shows high specificity, with a false positive rate lower than 0.1%. Finally, we have shown that using a naïve approach based on a frequency threshold as a cutoff to separate real mutations from errors results in extremely high false positive rates, demonstrating the value of our approach.

Originally, when observing the data in Fig. 1, as a first approximation it seemed likely to assume that mutations with a similar frequency would be mutations shared on the same genomes. Accordingly, we had hypothesized that at least two clusters of mutations (T1764-/G3114A/A1664G and A1611G/A1744G/T1440C/G1906A/A535G) would be present on the same genomes. This turned out to be only partially true: mutations with similar frequency were sometimes indeed on the same genomes (e.g., T1764-/G3114A), but sometimes completely not (the former two and A1664G) (Table 2). These results illustrate the utility of MinION to resolve the relationships among mutations, and its advantage for differentiating variants with mutations displaying similar frequencies.

We further used our approach to perform the reverse analysis: when analyzing the mRNA of the yeast gene enolase, our analysis suggested that the mRNA population sequenced was not homogenous. This was then precisely verified by Illumina sequencing of the same population. Remarkably, this analysis shows that (a) AssociVar can be used to analyze different types of data, ranging from virus genomes to mRNA of any organism, and (b) AssociVar can be used without sequencing a control sequence. We note that this requires more caution, since our analysis of MS2 showed that spurious associations between mutations may be created artifactually by the sequencing process itself. Use of AssociVar without a control sequence requires the user to specify the threshold of the normalized chi square statistic. As with all methods, the specificity of AssociVar comes at the cost of sensitivity, and vice versa (Fig. 6). Nevertheless, it seems the best strategy we can suggest is to use a very high threshold, which is extremely effective for variants at a frequency higher than 5 or 10%.

It is important to delineate the limitations of our approach. For one, we cannot distinguish haplotypes/strains that differ from each other at one position only, because our method relies on the association between two positions containing real mutations. Similarly, if two strains differ at very proximal loci, AssociVar will also fail, since we filter out associations between mutations that are less than 15 base pairs apart. We postulate that the presumably artefactual associations we observed between proximal loci are induced by the RNA (or DNA) passing through the pore of the sequencer. Finally, we also noted specific patterns of mutations that were reproduced between our control sequence and the two experimental populations of MS2. This suggested that perhaps sequence context and/or RNA secondary structure also induce specific errors in MinION. These findings require further investigation.

Although our method is ideal for direct RNA or direct DNA sequencing, we also used the method for cDNA that was amplified from RNA in the case of the Zika virus analysis (*30*) (Fig. S4). When we tried to reconstruct the known haplotypes present in this sample, our method did not fully succeed to recapitulate the haplotypes (data not shown). One possible explanation for this is that during the amplification step, either chimeric sequences of both strains were created, or PCR recombination occurred, breaking down some of the linkage between sites. In such cases, the use of AssociVar is limited to the detection of mutations only, and this further suggests that direct RNA/DNA sequencing may be preferable.

To summarize, we anticipate that due to its ease of use and advantages listed above, direct long read sequencing using MinION will be increasingly valuable in the field of virus genetics and in additional diverse fields such as transcriptome studies, cancer genetics, and microbiology. The AssociVar approach we suggest herein is simple and applicable to any organism, and as such we hope it will be a useful addition to the genomics toolbox in multiple fields.

## Acknowledgements

We thank David Burstein and Tzachi Hagai for critical reading. This study was in part funded by a grant to AS from the Israeli Science Foundation 1333/16, by grants from the VW Foundation (Experiment! 94070) and BSF (2016671) to UG. UG is also supported by the European Research Council (grant ERC-AdG 787514).

