## Supplementary for "Direct Sequencing of RNA with MinION Nanopore: Detecting Mutations based on Associations"

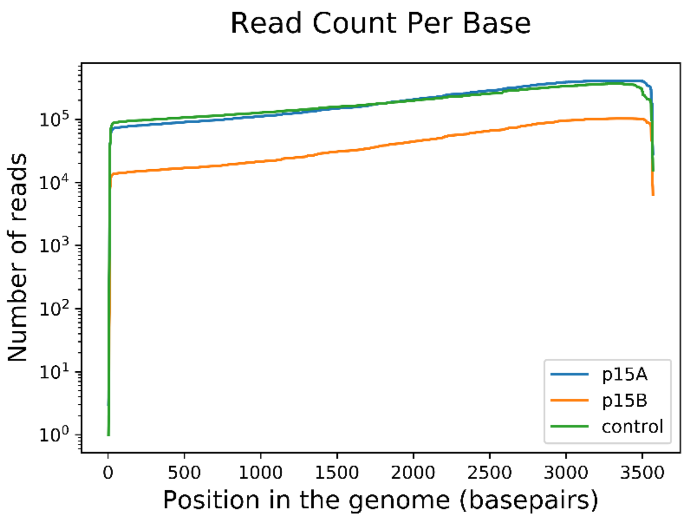


**Figure S1. Coverage plots of MinION direct RNA sequencing of MS2 in p15A, p15B and control.** The biased slope towards the left highlights the distribution of read lengths, where a large proportion of shorter reads was produced by MinION, and a smaller fraction reached the full length of MS2 (~ 3,569 bp).

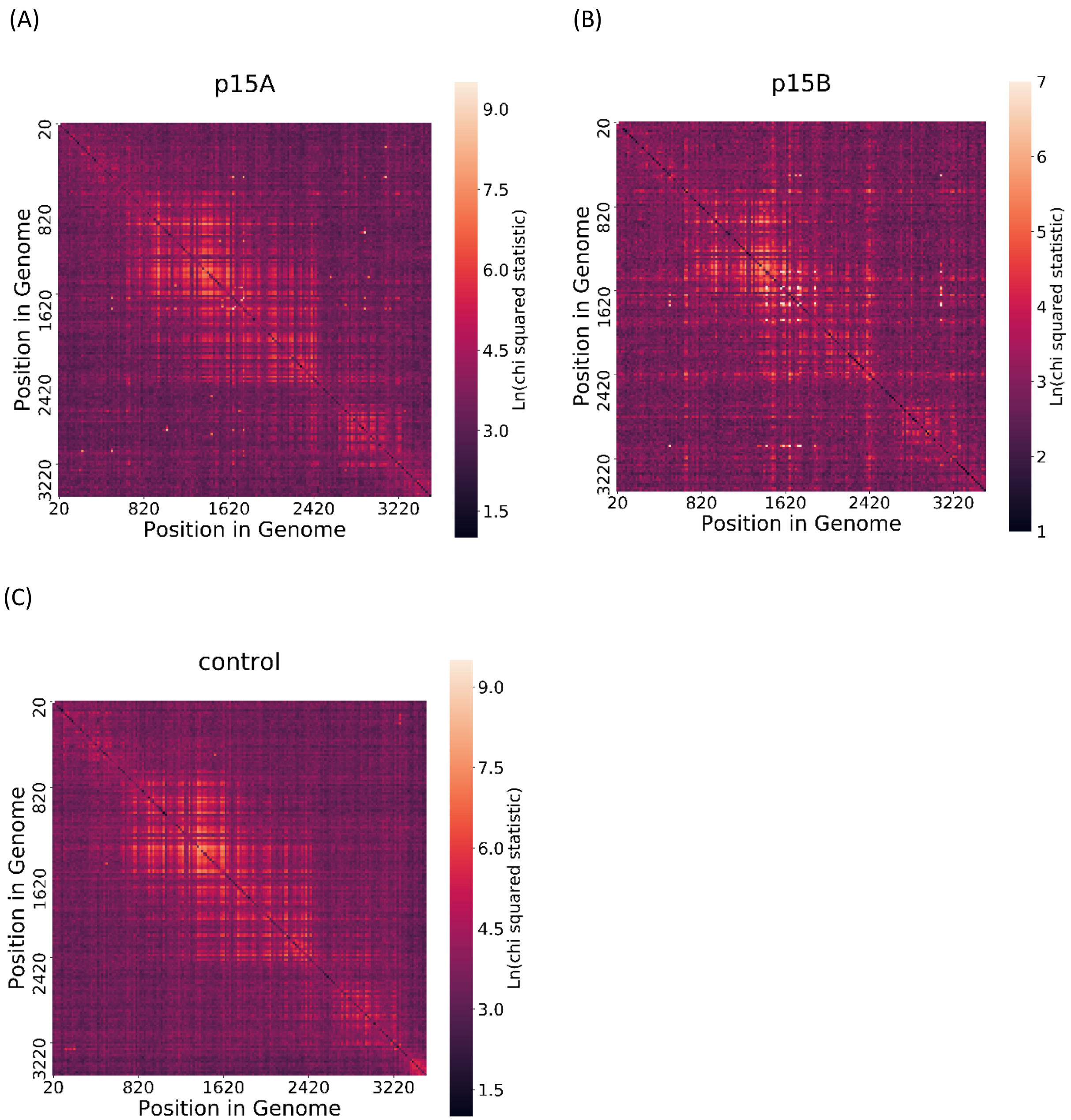


**Figure S2. Heatmap of chi square results.** The results are presented as the natural logarithm of the chi square statistic. (A), (B) and (C) show the results for p15A, p15B and control respectively. Results are scaled down: each group of 20x20 statistics are presented as the maximum statistic of the group. Statistics for two positions 15 base pairs or less away from each other were removed. The light vertical and horizontal lines show a wider distribution of chi-square scores for some positions. This seems to happen at positions with a variant frequency close to 50% and reflects a property of the chi-square test, but it is not a reflection of the probability of these positions having a real variant. Samples show an overall very similar pattern, meaning that MinION sequencing has a tendency towards specific errors for a specific genome sequenced. The relationships between two real mutations are seen as brighter than their surrounding area.

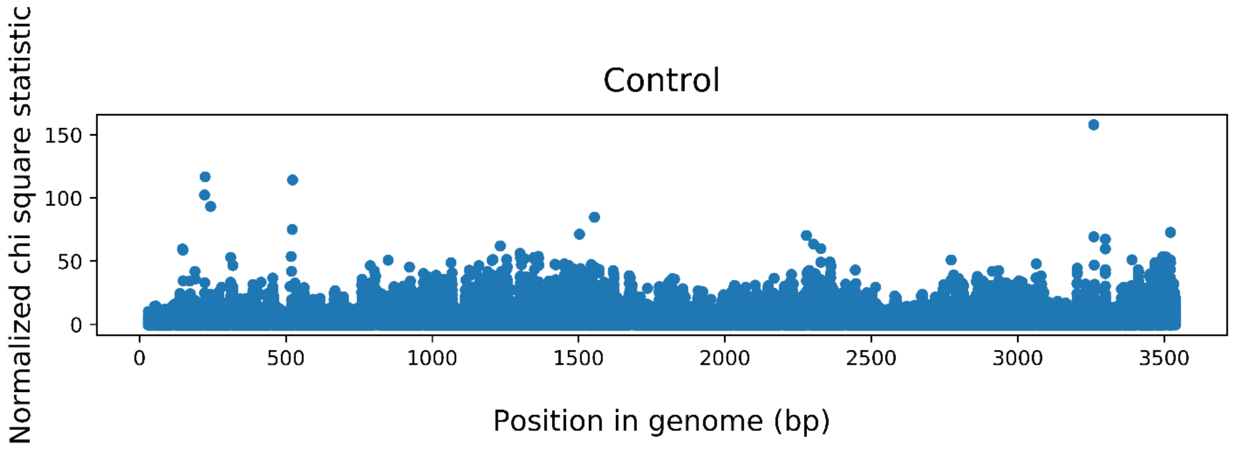


**Figure S3. Chi-square statistics plotted along the genome for the control sample minION sequencing, p1A**. Details are as in Fig. 4, yet notice the scale difference between the two plots. The control sample associations are much less prominent than the associations for p15A and p15B.

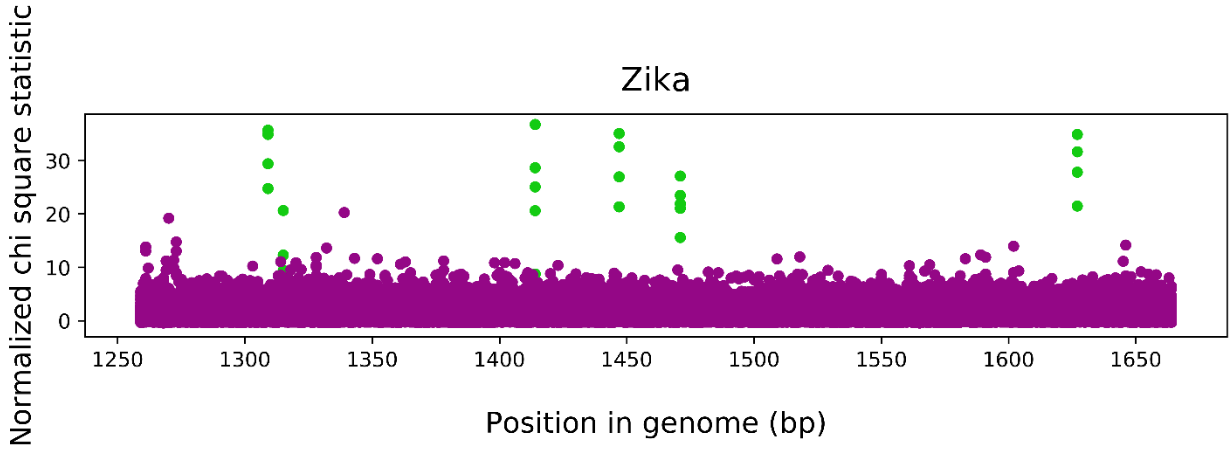


**Fig S4. Chi-square statistics plotted along the genome for the Zika virus amplicon of positions 1229-1665.** The association between the six positions with true mutations are marked in green. As can be seen, five out of the six mutations have associations higher than any associations between other positions, whereas the associations for the sixth mutation at position 1315 are slightly less significant.

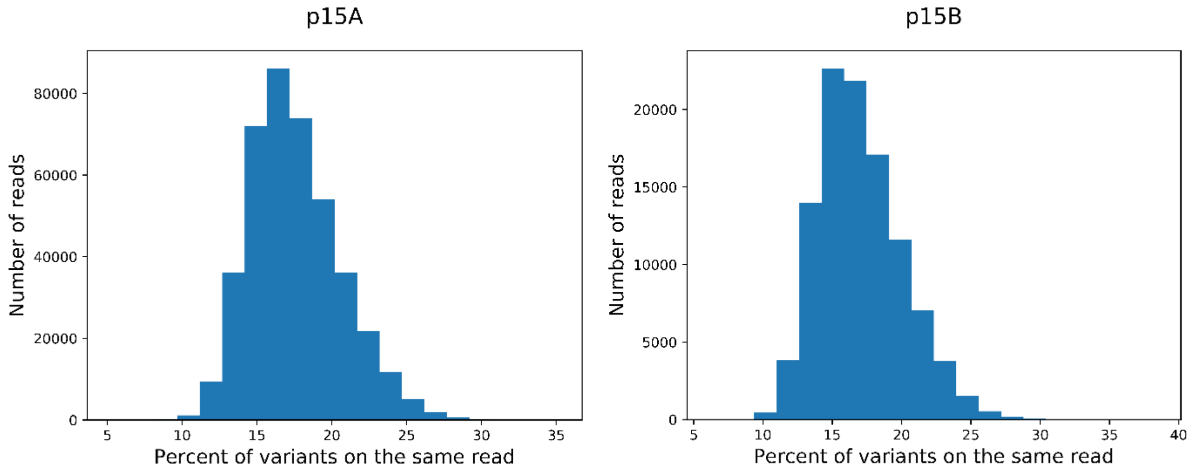


**Figure S5. The distribution of the percent of variants observed on the same MinION read**.

**Table S1. The differences between the MS2 reference (GenBank ID V00642.1) and the consensus sequence from passage 1.**

| Position | Passage 1 consensus | V00642.1 |
|---|---|---|
| 2002 | A | G |
| 2004 | C | G |
| 2005 | G | A |
| 2006 | G | T |
| 2007 | T | C |
| 2159 | C | T |
| 2160 | T | C |
| 2426 | C | T |
| 2429 | T | C |
| 2591 | T | A |
| 3038 | C | T |
| 3451 | C | - |
| 3452 | C | - |
| 3463.01 | - | C |
| 3463.02 | - | C |

**Table S2. Pairs of positions having a normalized chi score higher than the normalized chi-square cutoff of 114 defined by the control.** The results also include a "local maximum" column, determining whether the normalized chi score of the pair answers the condition of being higher than its surrounding, thus being considered a real mutation by our method. Positions that are identified as false positives when comparing our method to the Illumina results are marked in red.

| Position 1 | Position 2 | Chi score statistic | Normalized chi score statistic | Local maximum |
|---|---|---|---|---|
| **P15A** | | | | |
| 1050 | 2901 | 6194.947 | 641.6877 | TRUE |
| 1664 | 1764 | 9216.712 | 576.5063 | TRUE |
| 2901 | 1050 | 6194.947 | 512.5677 | TRUE |
| 1688 | 1744 | 3130.478 | 501.9177 | TRUE |
| 1560 | 1744 | 2547.602 | 460.6541 | TRUE |
| 531 | 3100 | 3206.283 | 438.3003 | TRUE |
| 1764 | 1664 | 9216.712 | 423.9477 | TRUE |
| 3100 | 531 | 3206.283 | 402.5826 | TRUE |
| 1744 | 1688 | 3130.478 | 387.3511 | TRUE |
| 3105 | 252 | 1675.803 | 376.9591 | TRUE |
| 535 | 1764 | 2385.867 | 367.655 | TRUE |
| 252 | 3105 | 1675.803 | 351.425 | TRUE |
| 535 | 1664 | 2103.958 | 324.203 | TRUE |

| | | | | |
|---|---|---|---|---|
| 1744 | 1560 | 2547.602 | 315.1994 | TRUE |
| 1663 | 1764 | 1605.107 | 235.4402 | FALSE |
| 1131 | 1764 | 1312.509 | 223.9578 | TRUE |
| 2953 | 1475 | 1096.916 | 221.1228 | TRUE |
| 2585 | 1475 | 1166.538 | 215.7766 | TRUE |
| 1051 | 2901 | 1367.668 | 209.2721 | FALSE |
| 1549 | 1131 | 1076.928 | 200.0303 | TRUE |
| 1131 | 1549 | 1076.928 | 183.7197 | TRUE |
| 2735 | 1724 | 615.3027 | 151.5377 | TRUE |
| 2901 | 1764 | 1801.675 | 149.0306 | TRUE |
| 1763 | 1664 | 909.4032 | 142.3082 | FALSE |
| 1475 | 2585 | 1166.538 | 142.1301 | TRUE |
| 1475 | 2953 | 1096.916 | 133.6314 | TRUE |
| 1664 | 535 | 2103.958 | 131.567 | TRUE |
| 2356 | 1764 | 539.1442 | 126.3444 | TRUE |
| 1655 | 1764 | 475.5166 | 119.5437 | TRUE |
| **P15B** | | | | |
| 3114 | 1764 | 4009.217 | 361.0727 | TRUE |
| 1440 | 1744 | 5119.376 | 333.4319 | TRUE |
| 1611 | 1744 | 5385.559 | 332.46 | TRUE |
| 1764 | 3114 | 4009.217 | 318.1413 | TRUE |
| 1744 | 1611 | 5385.559 | 302.9105 | TRUE |
| 1744 | 1440 | 5119.376 | 287.9376 | TRUE |
| 1440 | 1611 | 4388.441 | 285.8182 | TRUE |
| 1611 | 1440 | 4388.441 | 270.8996 | TRUE |
| 3113 | 1764 | 980.8612 | 256.0495 | FALSE |
| 3112 | 1764 | 1038.127 | 253.3373 | FALSE |
| 3114 | 1664 | 2803.999 | 252.5169 | TRUE |
| 1441 | 1744 | 778.0159 | 239.5114 | FALSE |
| 1664 | 3114 | 2803.999 | 229.2336 | TRUE |
| 1664 | 1764 | 2671.664 | 218.4126 | TRUE |
| 1906 | 1744 | 1782.185 | 218.3015 | TRUE |
| 1764 | 1664 | 2671.664 | 211.9908 | TRUE |
| 535 | 3114 | 705.8766 | 205.9131 | TRUE |
| 1906 | 1611 | 1670.611 | 204.6286 | TRUE |
| 1441 | 1611 | 644.9021 | 198.5045 | FALSE |
| 1593 | 1744 | 1016.355 | 197.3718 | TRUE |
| 1593 | 1611 | 1006.947 | 195.5436 | TRUE |
| 1730 | 535 | 300.2778 | 188.1661 | TRUE |
| 3113 | 1664 | 715.8328 | 186.8283 | FALSE |
| 1906 | 1440 | 1505.701 | 184.4198 | TRUE |
| 3112 | 1664 | 745.9047 | 181.9814 | FALSE |
| 535 | 1764 | 614.4724 | 179.2326 | TRUE |

| | | | | |
|---|---|---|---|---|
| 1593 | 1440 | 892.3381 | 173.2721 | TRUE |
| 1592 | 1611 | 822.6934 | 152.2253 | FALSE |
| 1592 | 1744 | 815.9222 | 150.971 | FALSE |
| 1592 | 1440 | 751.9454 | 139.1195 | FALSE |
| 3109 | 1764 | 361.04 | 127.5835 | TRUE |
| 1763 | 3114 | 318.8337 | 119 | FALSE |

### Supplementary text

#### Haplotype/Strain Identification Analysis

A. For variants X, Y and Z, find all possible combinations of variants (haplotypes) that were observed in the sequencing data.

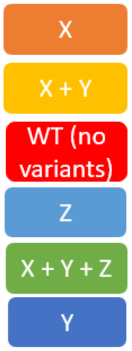


B. Create a group for every base variant, where each group contains all of the haplotypes that contain that variant. WT is also treated as a base for a group which will include all of the observed haplotypes. Haplotypes will hence be present in more than one group.

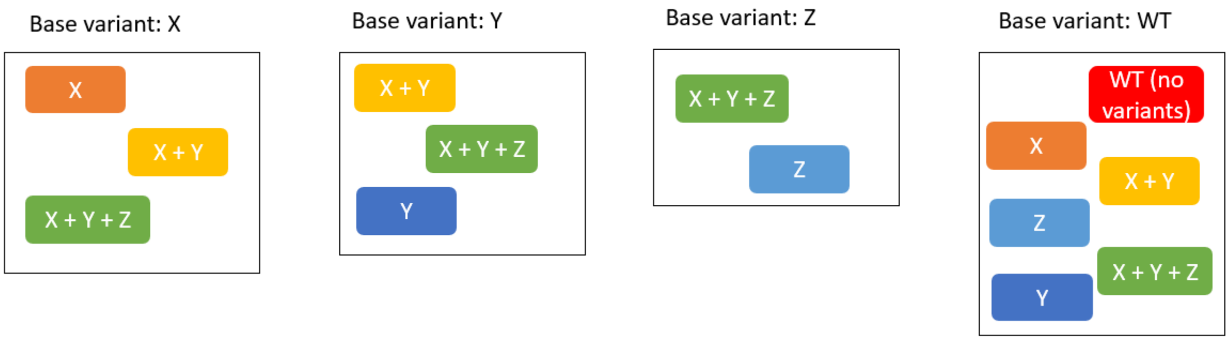

C. Within group, sort by the absolute frequency of the haplotype in the sample, and also calculate the relative frequency of each haplotype in its base variant group.

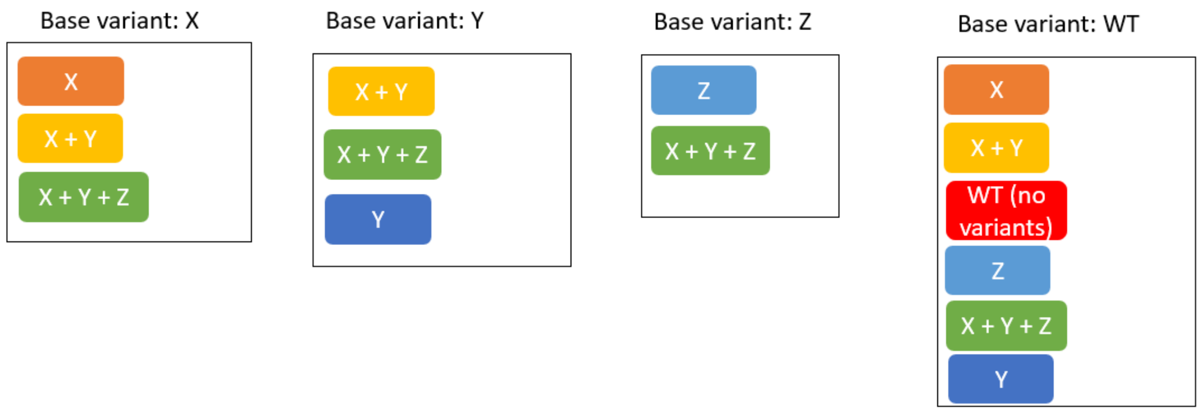


D. Within each group, iterate through the haplotypes from highest frequency to lowest, classifying each haplotype as reliable or not. The first haplotype is automatically classified as reliable. For every following haplotype, we compare its relative frequency with the probability that it is created by technical errors from the closest haplotype classified as reliable, called its parent haplotype, using the inferred error threshold. For example, if a haplotype has an additional deletion and substitution when compared to its parent haplotype, we require that its relative frequency be higher than the product of 0.214x0.237=0.051 to be classified as reliable (using the 95$^{th}$ percentile error frequencies from Table 1). For example, for base variant X:

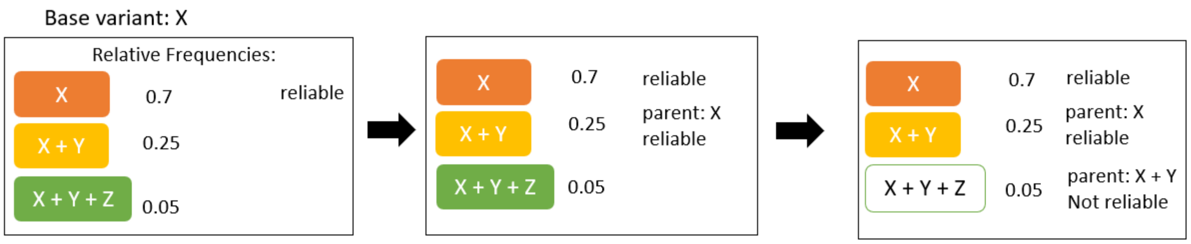


We do this within every group:

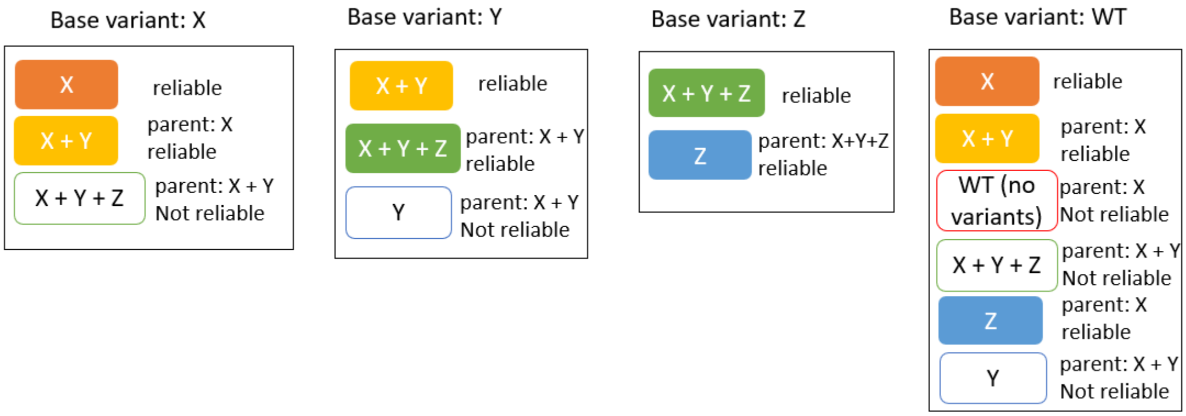


E. For haplotypes appearing in more than one group, it is enough to be classified as reliable in one group to be classified as reliable overall. Finally, we report the overall list of reliable haplotypes and recalculate their relative frequency within the set of reliable haplotypes.

Reliable strains:

| | X | X + Y | X + Y + Z | Z |
|---|---|---|---|---|
| Absolute frequencies: | 0.56 | 0.2 | 0.04 | 0.03 |
| Recalculate frequencies within reliable haplotypes: | 0.67 | 0.24 | 0.05 | 0.04 |
